# *Treponema pallidum* TprD and TprK Are Adhesins and Promote Spirochetes Opsonophagocytosis

**DOI:** 10.64898/2026.01.04.697605

**Authors:** Kashaf Zafar, Onyedikachi C. Azuama, Linda Xu, Lorenzo Giacani, Nikhat Parveen

## Abstract

*Treponema pallidum* subspecies *pallidum* (*T. pallidum*) causes systemic syphilis, exclusively infects humans in nature and can persist for decades in the absence of treatment despite generating robust adaptive immune responses. The *<u>T</u>. <u>p</u>allidum* <u>r</u>epeat (Tpr) family of outer membrane proteins are immunogenic but have been implicated in immune evasion due to antigenic variation, indicating that Tprs are virulence factors displayed on the spirochetes surface. Long-term survival of *T. pallidum* is largely attributed to the sparse surface-exposed outer membrane proteins and antigenic variation exhibited by the major surface protein TprK, which undergoes phase variation. Mechanism of antigenic variation has been studied for decades; however, functions of Tprs in this extracellular pathogen have not been experimentally determined. In this study, we determined TprD and TprK location, their role in adherence and in clearance of *T. pallidum* by macrophages. Using our now established heterologous surrogate system and gain-in-function approach using non-adherent related spirochete, the B314 strain of *Borrelia burgdorferi*, we show that both TprD and TprK are surface exposed to some extent on engineered *B. burgdorferi* as well as on *T. pallidum* Nichols and SS14 strains. We further demonstrate that both proteins mediate adherence to different mammalian cells *in vitro* and mouse antibodies generated against TprD and TprK putative surface loops bind spirochetes and promote J774A.1 macrophages-mediated opsonophagocytosis. Thus, surface-exposed adhesins TprD and TprK of *T. pallidum* contribute to binding to different types of cells which likely reflect the pathogen’s ability to colonize different tissues, and they are also targets of opsonic antibodies.

**IMPORTANCE:** Syphilis remains a major global public health challenge and is exacerbated by the rising number of cases of congenital infection and increased risk of HIV acquisition and transmission in syphilitic patients. A critical barrier to improving understanding of the molecular basis of syphilis pathogenesis owe to fragility of *T. pallidum*, inability to grow it in pure culture, and difficulty in generating knockout mutants in predicted virulence factors due to their possible essential role in spirochetes viability. Our findings provide experimental evidence linking Tpr proteins to host cell adherence, their ability to generate humoral immune response in the rabbit model of infection as well as in humans which could facilitate clearance by macrophages. Demonstration of TprD and TprK as the targets of opsonic antibodies here highlight their potential as protective immunogens and emphasizes importance of their inclusion in the cocktail to produce effective vaccine against syphilis.

## INTRODUCTION

*Treponema pallidum* subspecies *pallidum* (*T. pallidum*) spirochetes are the causative agent of syphilis, which is a re-emerging global health problem characterized by systemic infection and lifelong disease in the absence of treatment. The World Health Organization (WHO) estimates that 5.6-11 million new syphilis infection cases among men and women aged 15–49 years annually, and a prevalence of 18-36 million cases globally (1–5). In addition, congenital syphilis (CS), caused by transplacental transmission of *T. pallidum* during pregnancy, has been increasing around the world including in the Western countries. Vertical transmission of *T. pallidum* is a significant cause of stillbirth, neonatal death, and CS in newborns associated with severe neurologic and developmental defects with estimated ∼700,000 cases reported per year (6–14). The increase in CS to 3,882 cases in the United States in 2023 and a further 2% surge in 2024 underscores the shortcomings in disease control even in high-income nations (15–17). Increased HIV transmission by 2-5 times in symptomatic syphilis patients has been reported (18, 19), further making *T. pallidum* infection a global public health problem.

The genome of *T. pallidum* is of relatively small size of ∼1.14Mbp, which encodes >1000 proteins (20). In nature, it is an exclusively human pathogen. *T. pallidum* pathogenesis and persistence is attributed mainly to the architecture of its outer membrane, with few and poorly expressed surface-exposed proteins, , antigenic variation in TprK, and stochastic expression of some surface proteins (21–25). By limiting the density of its surface antigens, this pathogen becomes difficult for the host immune system to detect and eliminate from host (26, 27). The characterization of the molecular basis of host- *T. pallidum* interactions and the understanding of syphilis pathogenesis remain quite challenging due to the fragility of spirochete outer membrane, inability to cultivate it continuously in pure culture *in vitro* and difficulty in generating mutants (28, 29) albeit there have been some successful attempts recently in inserting antibiotic cassettes and reporters in *T. pallidum* genome (3). Therefore, the functional assessment of *T. pallidum* virulence factors currently has been limited to the application of recombinant proteins and surrogate systems (25, 30–36).

In contrast to *T. pallidum*, the structurally and physiologically related Lyme disease causing spirochete, *Borrelia burgdorferi* with a genome size of ∼ 1.52 Mb expresses approximately 132 surface lipoproteins, which include known outer surface proteins (Osps) and known virulence factors. The majority of surface proteins of *B. burgdorferi* are highly immunogenic and dynamically regulated in response to colonization and survival in diverse environments (e.g., tick vs. mammalian host) and are encoded mostly by genes located on endogenous circular and linear plasmids that can get lost during long-term *in vitro* cultivation rendering spirochetes no-infectious (37). To overcome the challenges that hinder direct functional studies with physiologically active *T. pallidum,* we previously developed a heterologous expression system that enables the assessment of the subcellular localization of *T. pallidum* proteins and allow evaluation of their functions. We have successfully used a non-infectious and non-adherent derivative of the *B. burgdorferi* B31 strain, B314, as a surrogate to examine the role of some *T. pallidum* proteins (25, 34, 38–40). B314 is a highly passaged strain that has lost various endogenous plasmids and virulence functions, thus is useful as a heterologous expression system to investigate adherence mechanisms of other pathogenic spirochetes. This gain-of-function approach has contributed to progress towards elucidating structural components of *T. pallidum,* in characterizing the function of some *T. pallidum* surface proteins and validated their potential as vaccine candidates (40–42). Although the mechanisms of *T. pallidum* persistence have not been completely elucidated, stochastic expression of *T. pallidum* proteins on surface and antigenic variation of prominent TprK protein by phase variation likely facilitate survival despite stimulation of the specific adaptive immune response during infection (25, 43).

*Treponema pallidum repeat* (Tpr) family includes 12 proteins, some of which are known to be surface-exposed (20, 44, 45). They are predicted to possess β-barrel structures with surface loops exposed to the host immune system and have been regarded as key players in bacterial pathogenesis. They are categorized into three subfamilies according to amino acid homology: subfamilies I and II contain TprC, D, F, I and TprE, G, J, respectively, possess conserved amino and carboxyl terminal sequences but variable central amino acid sequences, whereas the subfamilies III (TprA, B, H, L, K) possess conserved sequences and scattered variable regions across their length (43, 44). These proteins exhibit homology to the major sheath protein (Msp) proteins of *T. denticola,* which have been previously implicated in cell attachment and also function as porins (44, 46, 47). TprK (encoded by the *tp0897* gene) is a highly expressed surface protein with a putative, cleavable signal peptide, predicted molecular weight of 52.7 kDa and has received considerable attention due to its allelic variation and the perceived role in immune evasion despite generating opsonic antibodies; however, surface exposure of this protein was previously challenged by one research group (44, 48, 49). TprK harbours seven discrete variable regions (V1-V7) that are predicted to form external loop at the host-pathogen interface. Furthermore, previous studies have shown that TprK elicits strong T-cell and antibody responses in the rabbit infection model (35, 44, 50–52).

The role of TprC/D in syphilis infection has also been emphasized by several researchers and these homologous proteins have been shown to be targets of humoral and cellular immune responses (43, 50, 53, 54). The TprC/D (Tp0117/131) are duplicated in the Nichols strain and has outer membrane localization and has been predicted to function as porins (20, 55–57). Although TprC and TprD are closely related, they exhibit differences in regions that may be exposed on the bacterial surface. TprC shows fewer hypervariable regions and is more conserved among different strains; however, TprD/TprD2 exhibit sequence and antigenic variation among strains (for example, in Nichols vs. SS14 strain), particularly in the external loops. The *tprD2* allele (found in the most currently circulating strains) possesses a 330 bp central variable region (CVR) and three smaller regions towards the end of the ORF that distinguishes it from *tprD*, which is found in the *T. pallidum* reference Nichols, Chicago, and Bal73-1 strains (43, 53, 58).

Additionally, the *tprC* locus in Bal3, MexicoA, Sea81-4, and 9 encodes a TprC protein with fewer amino acid variations compared to the TprC of the referenced Nichols, Chicago, and Bal73-1 strain (57). It is important to note that the sequence variations in TprD/C are localized in discrete variable regions. TprC/D paralogs have been reported as strong vaccine candidates in previous studies, where it was shown that the N-terminal conserved region induced strong T-cell and antibody responses during infection. Furthermore, immunization with this specific region of proteins reduced the development of early syphilis lesions and provided protective effects against *T. pallidum* infection in rabbits on challenge of vaccinated animals (53, 58).

For an extracellular pathogen like *T. pallidum,* receptor-mediated adherence to host tissues plays a crucial role in organs colonization and its long-term persistence in hosts. Upon infection, phagocytosis of opsonized treponemes by macrophages aid in the clearance of spirochetes from early cutaneous lesions (59–63); however, this clearance mechanism is rather slow, likely due to limited surface exposure of proteins. The N-terminal region is conserved among all members of subfamily I Tprs, eliciting antibody and T-cell responses. Immunization with this region was reported to minimize the development of syphilitic lesions upon infectious challenge (44, 53). Although TprD exists in two allelic forms: (TprD and TprD2), variation in these proteins do not occur during active infection. In contrast, TprK undergoes extensive intrastrain antigenic variation during infection (24, 64), via a mechanism based on non-reciprocal gene conversion from a pool of donor sequences. Furthermore, infection-induced antibodies target the V regions of TprK, with sequence diversity localized within the discrete variable regions, rather than the conserved regions. Previous investigation showed that the convalescent sera recovered from rabbits immunized with recombinant TprK recognize both variable and conserved regions (52).

Although TprD and TprK proteins have been studied for decades, the function of these proteins as adhesins remains undiscovered, albeit TprK was proposed to be a monomeric porins (65). We carried out this investigation to experimentally show that TprD and TprK proteins are also adhesins like their homolog, Msp of *T. denticola*. We utilized *B. burgdorferi* non-adherent B314 to express TprD and TprK proteins and demonstrate their adherence function shown by gain in B314 strain ability to bind to different mammalian cell lines. Furthermore, we observed that these proteins are expressed on the surface of engineered *B. burgdorferi* B314 and *T. pallidum* Nichols and SS14 strains, as well as are targeted by the specific antibodies to elicit opsonophagocytosis by macrophages *in vitro*.

## RESULTS

### Antibody production against TprD/TprK

The protein sequences of TprC/D (Fig. 1A) and TprK (Fig. 1B) chimeras were generated by GenScript in pET30a expression vector and confirmed by sequencing. The recombinant proteins purified from *E. coli* were used for Balb/c mice immunization and for different experiments in this study. We also generated clones in pJSB175 plasmid vector containing full TprD and TprK encoding sequences together with flanking DNA sequences that include potential promoter and downstream sequences (Fig. S1). B314 strain transformed by these clones allowed TprD and TprK proteins expression under native promoter. The characterized spirochete clones and antibodies generated were then used conduct functional studies as described below.

**Fig 1.**
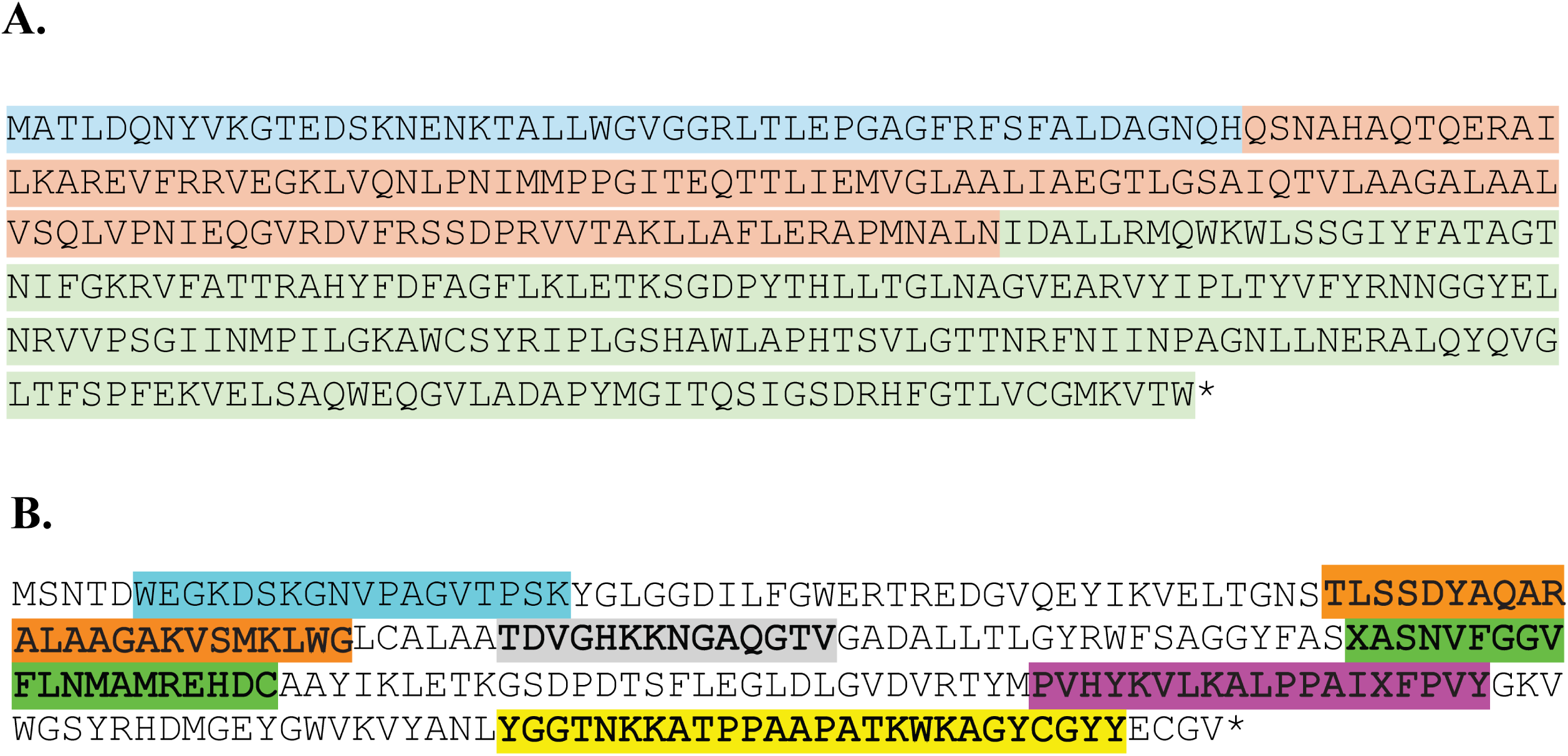
Sequence of Tpr chimera constructs generated by GenScript. DNA encoding TprC/D or TprK chimera was inserted in pET30a vector to express recombinant polyhistidine-tagged protein and purification by Ni-affinity chromatography. **(A)** Amino acid sequence of the synthetic TprC/D chimera. Previously identified MOSPN region is highlighted in blue, the CVR in orange, and the MOSPC β-barrel domain in green. **(B)** The chimera protein sequence for TprK includes variable outer epitopes with V2 highlighted in cyan, V3 in orange, V4 in gray, V5 in green, V6 in purple, and V7 in yellow color.

### Recognition of TprD and TprK on the Surface of *B. burgdorferi* by IFA using antibodies generated against recombinant proteins

We designated TprD and TprK expressing B314 strain as B314-TprD and B314-TprK, respectively, while the strain transformed with the pJSB175 vector alone is referred as B314-pJ and served as a negative control. Surface labeling of B314-TprD and B314-TprK with respective antibodies showed specificity of interaction because B314-pJ strain did not show any reactivity with these antibodies. (Figs 2D, 2L versus 2B, 2H). DNA staining by DAPI confirmed the presence of spirochetes in the respective fields. B314-TprD showed strong fluorescence signal with anti-TprD while punctate fluorescence was observed in B314-TprK treated with anti-TprK antibodies. All three strains (B314-pJ, B314-TprD, and B314-TprK) were also probed with antibodies against FlaB protein of *B. burgdorferi* under unpermeabilized condition in parallel. The lack of fluorescence indicates that the spirochete outer membrane integrity was maintained during this IFA. A strong TRITC signal detected in all strains after permeabilization validated FlaB antibodies recognition of periplasmic flagella (Fig. 2T, 2V and 2X). These results demonstrate that the antibodies raised against TprD and TprK specifically recognize their respective proteins on the surface of intact *B. burgdorferi.* This experiment was repeated four times.

**Fig 2.**
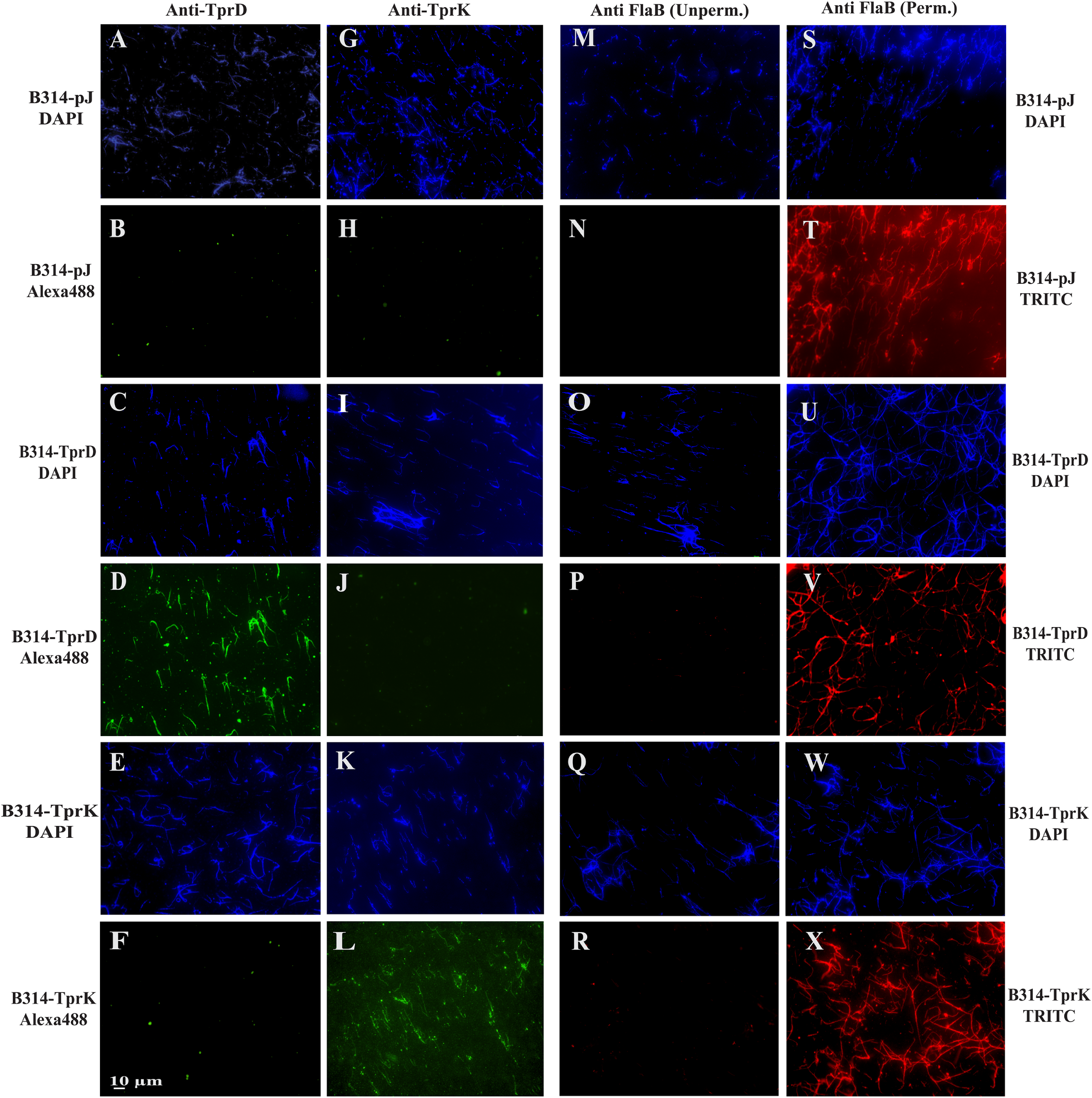
Detection of surface expression of TprD and TprK in *B. burgdorferi* B314 strain. IFA was performed to determine if TprD and TprK are located on spirochetes surface. To verify membrane integrity, anti-FlaB antibodies that recognize periplasmic proteins were used. Panels **A–F** show staining with anti-TprD antibodies, **G-L** with anti-TprK antibodies, both followed by AlexaFluor 488 conjugated secondary antibodies while panels **M-X** were treated with anti-FlaB antibodies followed by TRITC conjugated anti-mouse antibodies. B314 with the vector control, B314 (pJ), were labelled only with DNA dye with no detectable Alexa Fluor 488 fluorescence. B314-TprD exhibited strong surface labeling with anti-TprD while B314-TprK showed no detectable reaction with these antibodies. Similarly, B314-TprK strain was labelled with anti-TprK antibodies but not with anti-TprD antibodies. Panels **M–R** depict the lack of staining with anti-FlaB staining of unpermeabilized cells. Upon permeabilization, all strains treated with anti-FlaB antibodies displayed strong red fluorescence shown in panels **S–X**, confirming successful permeabilization and periplasmic localization of this flagellar protein. Scale bar depicts 10 μm.

### TprD and TprK on *B. burgdorferi* surface are recognized by secondary syphilis patient convalescent and infected rabbit sera

To further confirm that TprD and TprK are expressed in *B. burgdorferi* and are displayed on the surface IFA was performed using secondary syphilis patient serum (PS) and long-term (60 days of infection) infected rabbit serum (IRS). Again, staining by DAPI demonstrated the presence and distribution of spirochetes across all samples (Fig. 3A, 3C, 3E, and 3G). Surface labeling was observed when patient serum (Fig. 3B and 3D) or IRS (Fig. 3F and 3H) was used. Together, these results validate the expression of antigenic Tpr proteins and show that they are exposed on surface of the B314 strain.

**Fig 3.**
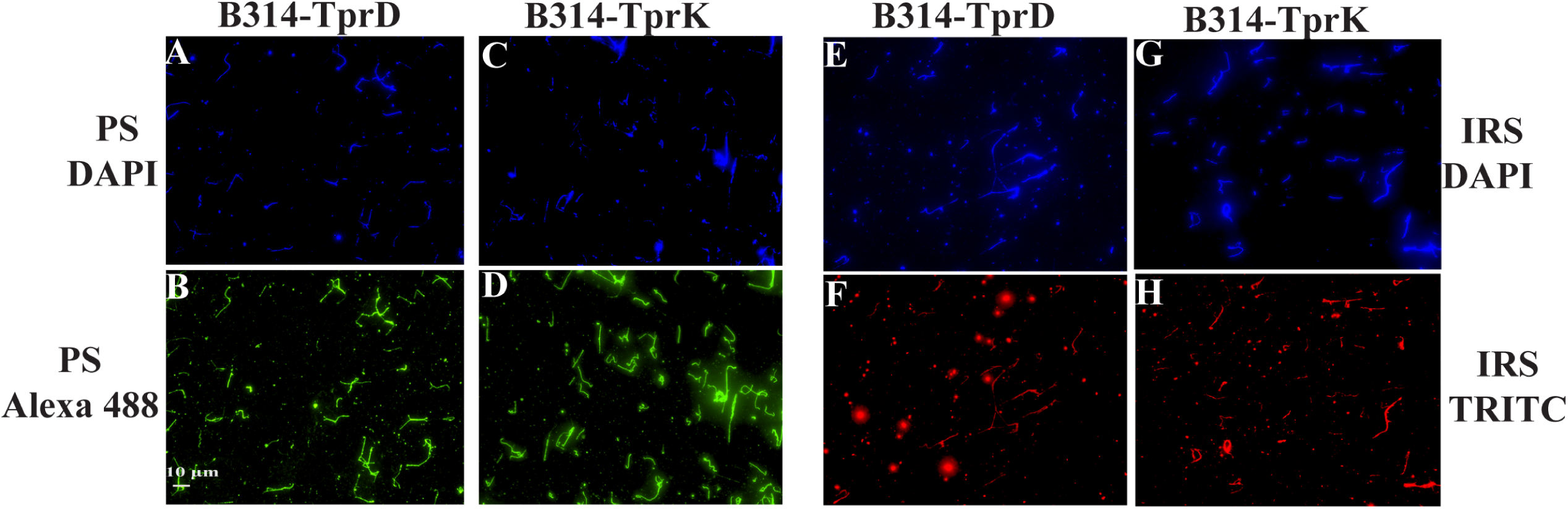
Immunofluorescence analysis using convalescent patient and rabbit sera confirming expression of TprD and TprK on *B. burgdorferi* surface. Panels **A, C, E and G** depict the presence of spirochetes in the fields by DAPI staining while spirochetes in **B and D** show robust Alexa Fluor 488 signal after secondary syphilis patient serum (PS)-treatment, demonstrating surface localization of respective proteins. (**F, H**) TRITC associated fluorescence shows surface staining after incubation with infected rabbit serum (IRS), confirming that TprD and TprK are surface-exposed. Bar indicates 10µm.

### Validation of mouse antibodies generated against chimeric TprD and TprK using *T. pallidum* Nichols and SS14 Strains

To ensure that antibodies we generated indeed recognize surface *T. pallidum* proteins, we conducted IFA with either Nichols, or SS14 strain co-cultured *in vitro* with Sf1Ep rabbit cells in the chambered slides. IFA revealed differential surface staining patterns for TprD and TprK on fixed Nichols and SS14 strains. Anti-TprK antibodies produced a clear, slightly punctate green fluorescence along the length of treponemes in both strains, indicating surface accessibility of TprK (Fig 4). Staining with anti-TprD produced scattered and punctate staining. Overall fluorescence signal was lower in *T. pallidum* compared to the surrogate B314 spirochetes expressing TprD and TprK (Fig 3 versus 4). Furthermore, relatively weaker staining of SS14 compared to that on Nichols strain could be due to differential surface expression and slight variation in the selected epitopes of TprD, which is duplicated in Nichols strain, and of TprK. As an internal control, unpermeabilized treponemes incubated with anti-FlaA antibodies showed no detectable fluorescence in either strain, consistent with FlaA being a periplasmic flagellar protein inaccessible for surface staining. After permeabilization, both Nichols and SS14 exhibited strong TRITC fluorescence, which confirms successful membrane disruption and validated antibodies reactivity to FlaA protein.

**Fig 4.**
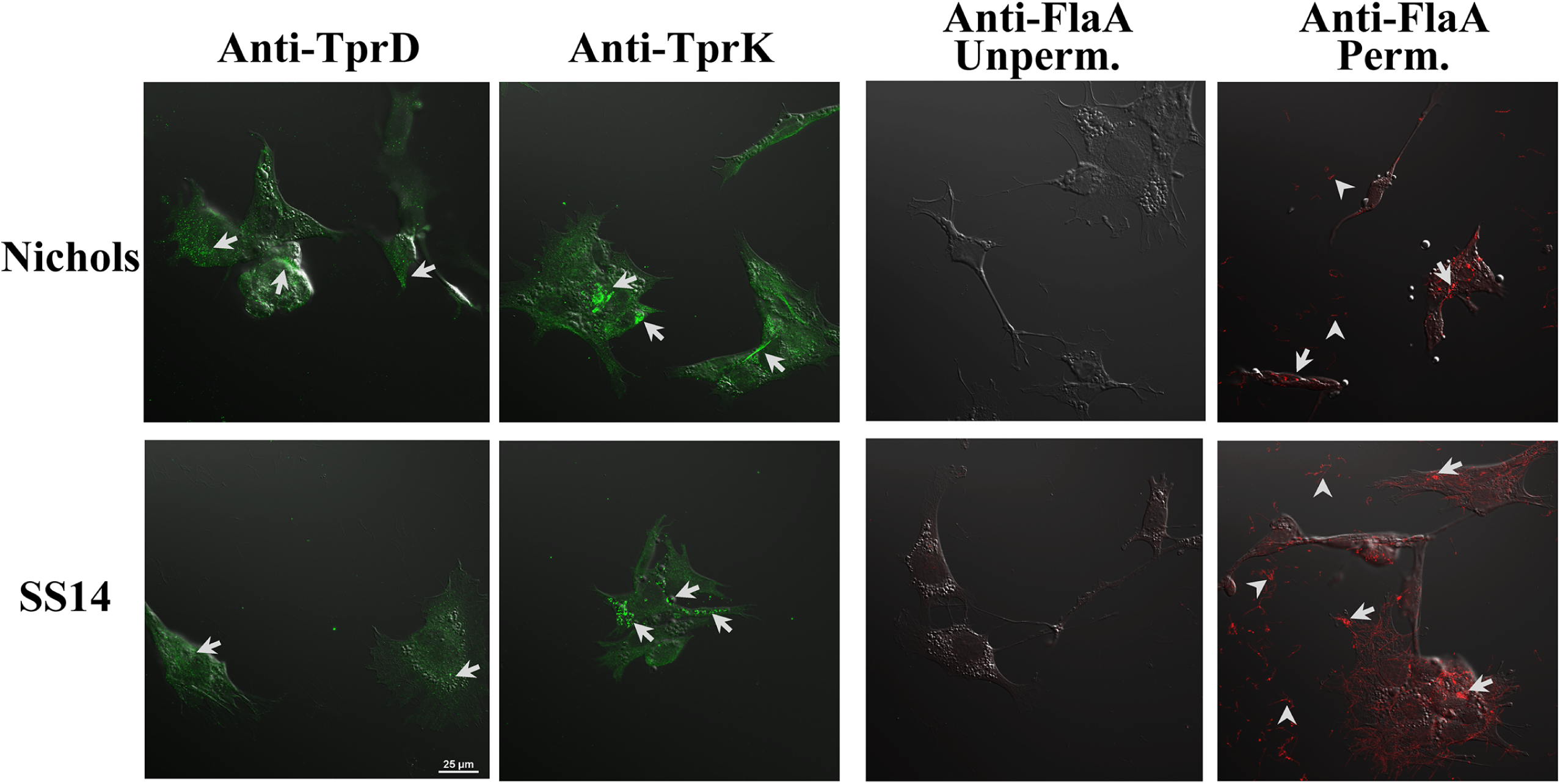
Detection of TprD and TprK on the surface of *T. pallidum* co-cultured with Sf1Ep cells *in vitro*. IFA conducted with co-cultured Nichols and SS14 strains by incubating with anti-TprK antibodies exhibited moderately punctate Alexa Fluor 488 staining along the cell surface, indicating surface accessibility of TprK. Anti-TprD staining produced discrete puncta suggesting limited recognition by our anti-TprD antibodies on *T. pallidum*. Unpermeabilized treponemes incubated with anti-FlaA antibodies showed no detectable signal because this is a periplasmic protein. Following permeabilization, both strains showed strong TRITC fluorescence after anti-FlaA treatment, validating reactivity of the primary antibodies. Bar depicts 20µm

### *T. pallidum* strain surface labels with secondary syphilis patient serum and infected rabbit serum

Nichols and SS14 strains labelling with a patient serum (PS) and infected rabbit serum (IRS) was detected by IFA revealing punctate green fluorescence for the syphilitic patient and red fluorescence for convalescent rabbit serum, confirming surface exposure of these proteins also in *T. pallidum*. Interestingly, the fluorescence using PS was more pronounced than that with IRS. SS14 strain showed higher signal with both primary polyclonal antibodies followed by respective fluorescent secondary antibodies (Fig 5).

**Fig 5.**
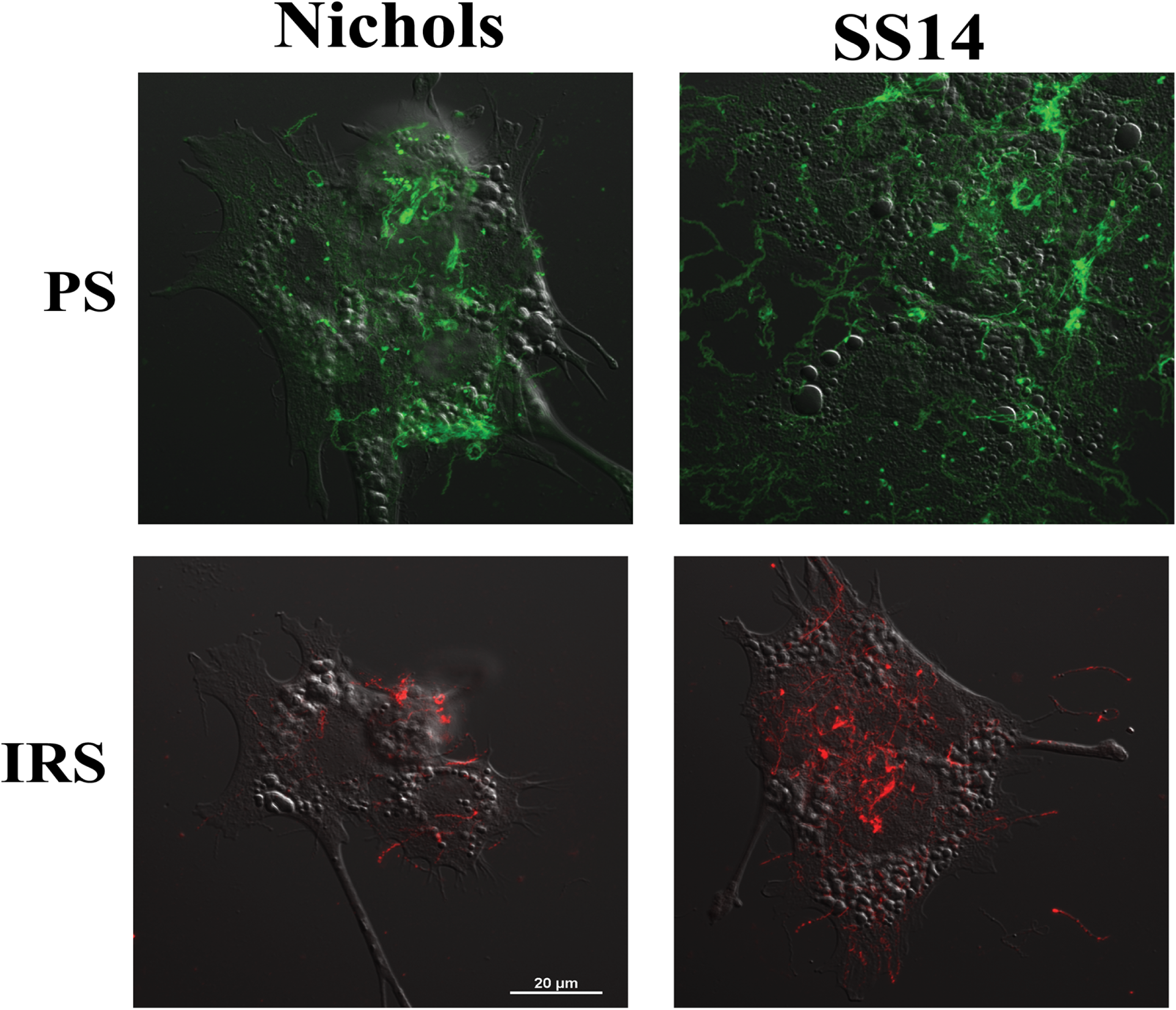
IFA-based detection of *T. pallidum* Nichols and SS14 strains surface proteins with secondary syphilis patient serum (PS) and infected rabbit serum (IRS). IFA with PS and IRS followed by FITC- and TRITC-conjugated secondary antibodies, respectively indicates low abundance of overall antigenic proteins on *T. pallidum* surface. Bar indicates 20μm size.

### Evaluation of Opsonophagocytosis

Since our antibodies were raised in mice, we used J774A.1 macrophage cell line for opsonophagocytosis experiment. To evaluate the contribution of TprD and TprK in antibody-mediated clearance, we first conducted opsonophagocytosis using the microscopy-based protocol that we previously optimized (Fig S2). Although some phagocytosis was observed after 3h of coincubation, results were ambiguous likely due to slow opsonophagocytosis documented for *T. pallidum* surface proteins takes longer as we previously showed with even with surrogate system expressing another *T. pallidum* protein where 6h coincubation was used (25). Therefore, we further employed a highly novel IncuCyte-based assay to ascertain opsonophagocytosis by J7741.1 cells

### Opsonophagocytosis of B314 strain expressing TprD and TprK by mouse macrophages

The IncuCyte system allows longer co-incubation of opsonized live bacteria with macrophages and has provision for automated data collection at selected timepoints. Thus, total red fluorescence generated after pH sensitive dye labeled bacteria enter the acidic phagosome is recorded. In this experiment, B314 strains were preincubated with pHrodo™ Red, succinimidyl ester together with respective anti-Tpr antibodies, and after washing added to J774A.1 cells in 24-well plate (with and without coverslips). Anti-OspC antibodies were included as a positive control and serum collected from uninfected, normal Balb/c mouse (NM) as negative controls.

Detectable fluorescence appeared after 4h incubation and continued to increase at later timepoints only when antisera were used and not with NM controls (Fig 6A). In fact, controls were almost indistinguishable from J774A.1 cell alone wells. Average of data collected from each well (nine points) without coverslip by machine at each time point are shown in this figure. After completion of the opsonophagocytic assay, coverslips from replicate wells were washed, mounted and imaged using Nikon ECLIPSE 90i fluorescence microscope using a 40x objective lens in the TRITC channel to observe phagocytosis throughout the coverslips (Fig 6B). Bright red fluorescence observed in most macrophages throughout indicate multiple red puncta confirming opsonophagocytosis data obtained by IncuCyte. Brightness of the phagosomes in images was more pronounced when multiple bacteria were simultaneously engulfed by a macrophage.

**Fig 6.**
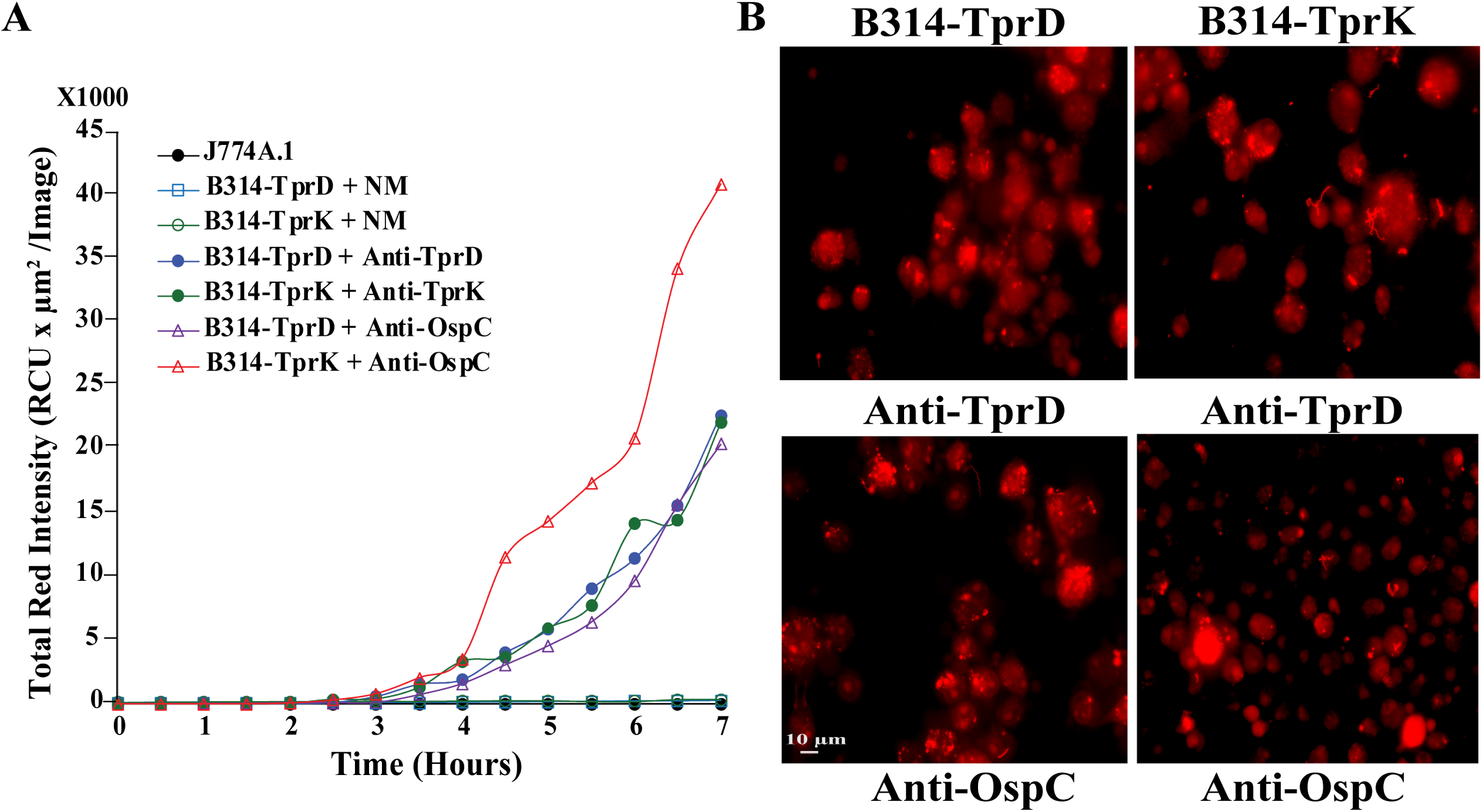
Opsonophagocytosis of *B. burgdorferi* B314 strain derivatives labelled with anti-TprD and anti-TprK antibodies using IncuCyte system. (A) Quantitative analysis of opsonophagocytosis showing increased albeit delayed but significant uptake of B314 expressing TprD or TprK when anti-TprD or TprK antibodies were used. *B. burgdorferi* surface lipoprotein OspC with antibodies against OspC were used as positive control and uninfected Balb/c normal mouse serum (NM) as negative controls. **(B)** Microscopic images showing a visual representation of Opsonophagocytosis after ∼16h of incubation in the IncuCyte machine.

### Opsonophagocytosis of *T. pallidum* strains after treatment with Anti-TprD and Anti-TprK antibodies

Successful outcomes of opsonophagocytosis of *B. burgdorferi* surrogate system incentivized us to conduct a similar assay with *T. pallidum* strains. Opsonophagocytosis of live *T. pallidum* Nichols and SS14 strains as indicated by red fluorescence signal, when reacted with our anti-TprD and anti-TprK antibodies, was relatively lower (Fig 7A) than that observed for B314 strain expressing these Tprs (Fig 6A). The most pronounced fluorescence signal was obtained with SS14 strain pretreated with anti-TprD antibodies. The imaging of multiple fields treated in parallel on coverslips in the same way in wells of the same plate visually demonstrate opsonophagocytosis of *T. pallidum.* The experiment was conducted several times and representative images from one experiment are shown (Fig 7B).

**Fig 7.**
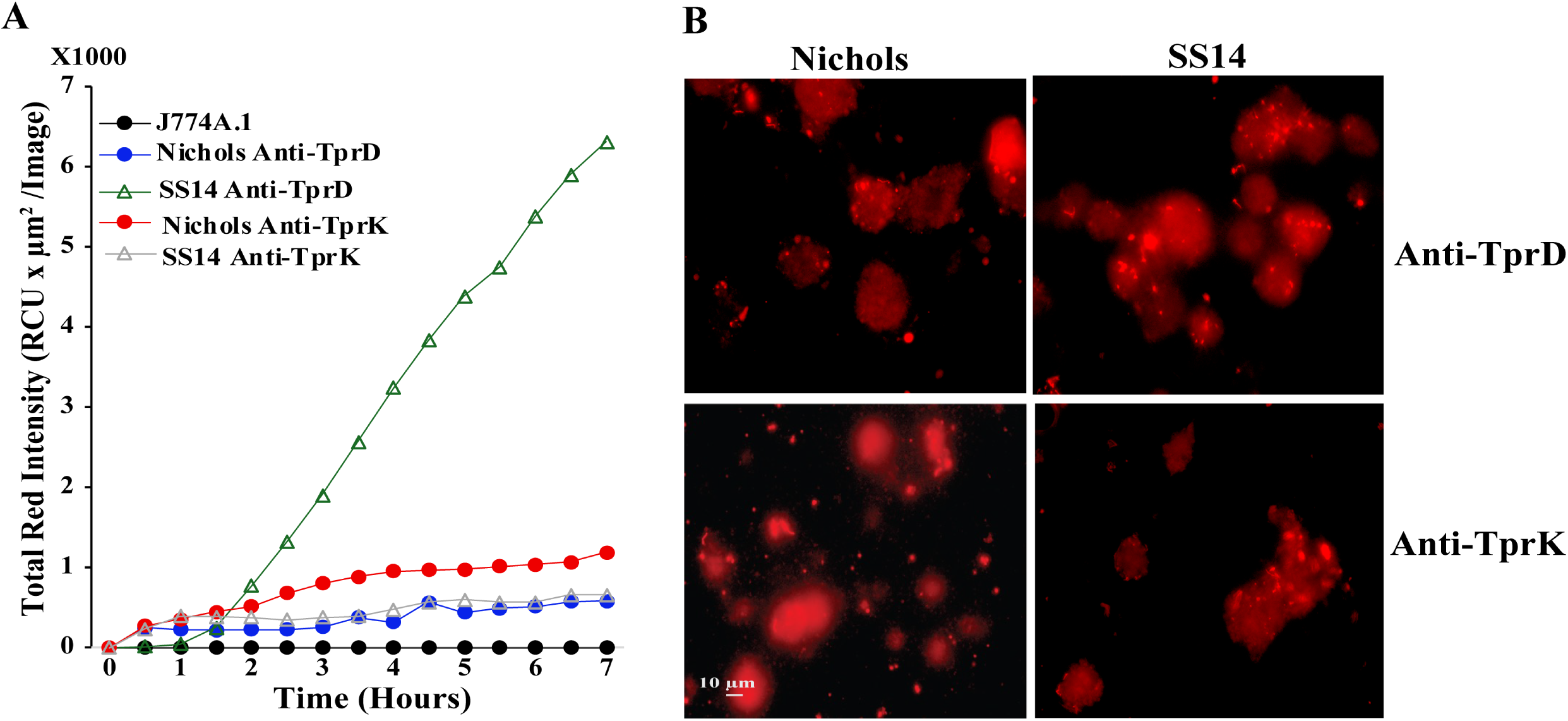
Opsonophagocytosis of *T. pallidum* Nichols and SS14 strains labelled with anti-TprD and anti-TprK antibodies. (A) Quantitative analysis of opsonophagocytosis showing increased uptake of Nichols and SS14 strains with prolonged incubation; however, most pronounced opsonophagocytosis was detected for SS14 treated with anti-TprK antibodies. (B) Microscopic images showing a visual representation of Opsonophagocytosis.

### B314 strain expressing either TprD or TprK gain ability to bind to 293, C6-glioma, and HeLa cell lines

We used radiolabeled B314 clones carrying either the shuttle vector alone (pJ), or the genes encoding TprD or TprK (Fig S1). B314 expressing Msp protein of *T. denticola* was included as a positive control. We conducted binding experiments with three different mammalian cell lines that are representatives of cell types possible targeted by *T. pallidum* during infection. Results from one representative experiment are shown here (Fig 8). In No cell control, all constructs showed only background levels of radioactivity associated with spirochete indicating no *B. burgdorferi* binding to the empty wells. B314 with the vector control exhibited minimal binding (∼1-3%) to all three cell lines while the expression of TprD or TprK significantly increased binding, reaching ∼7-9% for TprD and ∼9-11% for TprK, particularly on HeLa cells. The Msp-expressing clone demonstrated the highest level of binding with ∼12-13% on HEK293 cells, ∼17-18% on C6 cells, and ∼13-14% on HeLa cells (Fig 8A, 8B, and 8C). These results demonstrate that B314 acquires ability to adhere to host cells only when TprD, TprK, or Msp is expressed supporting roles of TprD and TprK of *T. pallidum* as adhesins.

**Fig 8.**
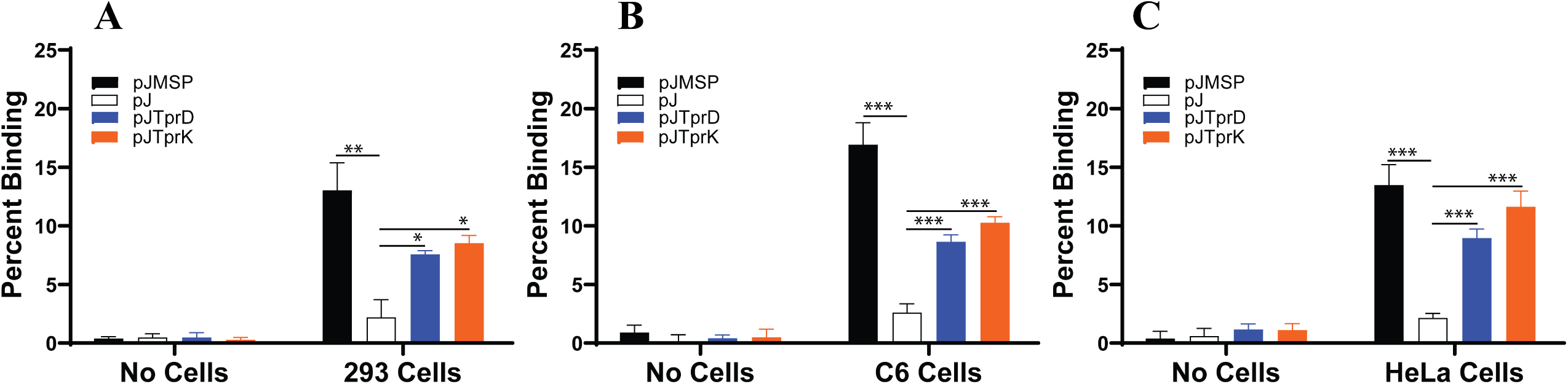
B314 acquires ability to bind to mammalian cells after TprD and TprK expression. Radiolabeled spirochetes carrying the shuttle vector pJSB175 alone (pJ), those expressing *T. pallidum tprD* or *tprK* genes or the *T. denticola* Msp encoding gene were incubated with 293, C6, or HeLa cell monolayers for 1h. After washing, no-cell controls yielded almost no detectable signal, the pJ vector control exhibited minimal binding while expression of TprD and TprK significantly increased binding. Msp, which produced the highest increase in binding. Bars represent mean ± SD from quadruplicate wells. Statistical analysis was conducted using a two-tailed unpaired student t test for unequal variance to determine significant difference between the paired groups and p values calculated (*p<0.05, **p <0.01, and *** p<0.001)

## DISCUSSION

The re-emergence of syphilis among young adults and the increase in cases of congenital syphilis are serious global health challenges that underscore the need for defining *T. pallidum* molecular pathogenesis mechanisms for development of effective diagnostic and preventative measures. Although some unique *T. pallidum* proteins are now recognized as its virulence factors, significant gaps remain regarding the subcellular location of various proteins, the transcriptional mechanisms of its virulence factors, and most importantly, functional assessment of its proteins that are expressed at significant levels. Electron microscopic examination and cryo-electron tomography previously showed molecular architecture of *T. pallidum* and scant presence of surface proteins (28, 66, 67). For extracellular pathogens like *T. pallidum* and *B. burgdorferi* that cause systemic diseases, receptor-mediated adherence to host cells is critical for organism persistence and long-term survival despite stimulation of a strong adaptive immune response.

The Tpr proteins have been evaluated by different researchers because of the presence of predicted N-terminal cleavable signal peptide and antigenic variation in TprK suggests them to be exposed on the *T. pallidum* outer surface and because of their potential contribution as virulence factors (43, 44, 49). Among these, TprK is the most extensively studied candidate (26, 48, 49, 60, 68). We decided to investigate the location of both TprK and TprD using robust immunofluorescence and opsonophagocytosis assays. Instead of using recombinant proteins that do not offer conformational integrity of outer membrane proteins, we employed a *B. burgdorferi* surrogate system and gain-of-function approach as we successfully used previously for other *T. pallidum* proteins (34, 36, 41). Specific surface labelling with antibodies generated against TprD and TprK putative surface epitopes chimeras only when TprD and TprK are expressed indicate that these are indeed surface proteins. Antigenic variation exhibited by TprK in *T. pallidum* and surface localization of the homologous Msp in *T. denticola* support our finding because this phenomenon is exclusively associated with surface-exposed proteins of microbes that cause long-term infection. Antigenic variation allows pathogens escape specific adaptive immune responses during infection. Furthermore, both Tpr proteins were recognized on the B314 strain surface by a secondary syphilis patient and a convalescent rabbit serum. Surface labelling also occurred on *T. pallidum* Nichols and SS14 strains grown in co-culture with rabbit epithelial Sf1Ep cells. In general, our results agree with previous demonstration that TprD/K proteins are surface exposed to some extent (44).

The outer surface molecules of extracellular pathogens, which are often immunogenic, play a major role in adherence and pathogenesis and in immune system mediated clearance because they are often targets of bactericidal or opsonic antibodies (44, 48). Previous studies have shown that antibodies produced in syphilitic serum against outer membrane proteins promote opsonophagocytosis (60, 62, 69, 70), and exhibit complement-dependent treponemicidal activity (71). Macrophage-mediated phagocytosis of opsonized *T. pallidum* is a crucial mechanism for their clearance from primary and secondary lesions (72–74), and opsonic antibodies are essential for killing treponemes by macrophages (59). Our immunolabeling and opsonophagocytosis experiments previously revealed that populations of *T. pallidum* are heterogeneous, consisting of antibody-binding and non-binding subpopulations (25, 48). Moreover, the organisms that bind to antibodies do so with slow kinetics (48, 72, 75). In this study, we used the IncuCyte system to examine opsonophagocytosis when spirochetes were pre-labelled with specific antibodies and a pH-sensitive dye. We found that anti-TprD and anti-TprK, and not NM control, promote opsonophagocytosis of both surrogate *B. burgdorferi* expressing either of these two proteins and two strains of *T. pallidum* by J774A.1 cells. Thus, our results agree with previous reports that showed that anti-TprK antibodies facilitate opsonophagocytosis of *T. pallidum* (44). These results also support our findings in which immunization with recombinant TprK or *B. burgdorferi-*expressing TprK induced partial protection in rabbits challenged with the *T. pallidum* Nichols strain (35, 44).

We further conducted functional analysis of TprD and TprK proteins. Since recombinant proteins may not accurately represent the surface epitopes of proteins, our related spirochete surrogate system represents adherence mechanism more closely to *T. pallidum*. Tprs are related to the Msp protein of *T. denticola,* which have been previously reported to be surface-exposed and exhibit functions both as adhesin and porin (44). Therefore, we included Msp as a positive control in our binding assays. Our results demonstrated that TprD/K expression confer ability to bind to the host cells by the B314 strain which is otherwise poorly adherent strain as demonstrated by binding of vector containing B314(pJ) strain (Fig 8). Our results suggest the role of TprD/K in tissue colonization during infection, potentially in skin lesions and/or during neurosyphilis manifestations (68, 76).

To summarize, our results emphasize the importance of surface-exposed Tpr antigens that generate opsonic antibodies as promising vaccine candidates. Since TprD/TprK are potent antigens, their ability to facilitate opsonophagocytosis support their inclusion in future vaccines to protect from infection by *T. pallidum*. In fact, testing of a multiple recombinant protein-based vaccine that combines the NH_2_-terminal fragment of TprK and the NH_2_-terminus fragments shared by Tpr Subfamily 1 proteins, together with Tp0751, is currently in progress in other laboratories (77). Recently, Molini and colleagues (58) observed cross-reactivity of antibodies towards variant peptides of TprC, TprD, and TprD2 in their quest to identify B-cell epitopes across TprC/D variants which also incentivizes exploration of these Tprs as vaccine components. Therefore, a multivalent vaccine that includes Tprs antigenic epitopes offer high promise of production of an effective and successful vaccine in the future.

## MATERIAL AND METHODS

### Ethics Statement

The *T. pallidum* Nichols and SS14 strains grown in *in vitro* co-culture system in the Giacani lab according to the Edmondson *et al.* protocol (78) in chambered slides were fixed with 4% paraformaldehyde made in PBS for IFA while those recovered from infected New Zealand White rabbits were used for opsonophagocytosis, both IFA and opsonophagocytosis experiments conducted in Parveen’s laboratory. All procedures followed the standards outlined in the Guide for the Care and Use of Laboratory Animals under the protocol reviewed and approved by the University of Washington Institutional Animal Care and Use Committee (IACUC protocol 4243–01; PI Lorenzo Giacani). Polyclonal antibodies against recombinant putative surface exposed loops of TprK and TprD were generated in BALB/c mice in Parveen’s laboratory using our previously established immunization procedure (79). These experiments were performed under Rutgers Biomedical and Health Sciences IACUC approved Parveen’s PROTO202000087 protocol. De-identified secondary syphilis patient serum used in this study was generously provided by the late Dr. Centurion-Lara.

### TprC/D and TprK putative surface epitopes chimeras used for antibody production

To generate polyclonal antibodies targeting the putative surface exposed domains of the TprC/D protein family identified previously (80), DNA constructs in pET30a vector to express recombinant chimera with defined Major Outer Sheath Protein N-terminal region (MOSP^N^), the Central Variable Region (CVR), and the C-terminal MOSP^C^ β-barrel domains were custom-designed and synthesized by GenScript (Fig. 1). We also designed clone in pET30a to produce chimeric protein that includes six variability domains of TprK (3). For Isopropyl β-D-1-thiogalactopyranoside (IPTG)-induced expression of each polyhistidine tagged protein, *E. coli* expression strain, BL21(DE3)pLysS was used and proteins purified using Nickel-affinity column using manufacturer’s instructions (Novagen). Purified proteins were subsequently used to immunize Balb/c mice to generate polyclonal antibodies as we described previously (81).

### Indirect Immunofluorescence Assay (IFA)

To determine TprD and TprK on *Borrelia burgdorferi* surface using B314 strain transformed with clones (Fig S1) generated in pJSB175 shuttle vector, IFA was performed. Spirochetes were harvested by centrifugation at 8,000 rpm for 10 minutes followed by three washes in PBS containing 0.2% BSA under identical centrifugation conditions. The pellet was then resuspended in PBS/0.2% BSA to achieve a concentration of approximately 2.5–5 × 10⁷/mL. Sterile 12-mm round coverslips were placed into 24-well plates, and each well was overlaid with 400 µL of HBSC buffer (25 mM HEPES, 150 mM NaCl, 1 mM MgCl₂, 1 mM MnCl₂, and 0.25 mM CaCl₂, pH 7.8) supplemented with 0.2% BSA and 0.1% glucose, followed by the addition of 100 µL of spirochete suspension. Plates were centrifuged at 2,000 rpm for 10 minutes at room temperature to promote organisms attachment to the coverslips. After removal of supernatants, spirochetes were fixed with 3% paraformaldehyde for 1h at room temperature and washed three times with PBS. Both unpermeabilized and methanol permeabilized (kept at −20°C) spirochetes were blocked with PBS containing 5% BSA and 5% heat-inactivated goat serum (blocking buffer) for 1h. Polyclonal antibodies generated against TprC/D or TprK surface epitopes were used at 1:1000 dilution with anti-FlaB monoclonal antibodies against periplasmic protein used as control to ensure integrity of outer membrane. Plates were incubated at room temperature for 2h with gentle rocking. After washing with PBS, all wells were permeabilized again with cold methanol for 10 minutes for DAPI staining (10µg/ml final) together with Alexa Fluor 488 or TRITC-conjugated secondary antibodies diluted at 1:100 for 1 hour at 37 °C in the dark and then washed. Coverslips were mounted onto glass slides and examined using Plan Apo λ objective in Nikon ECLIPSE 90i microscope. Similarly, IFA was conducted with *T. pallidum* strains grown in co-culture system in chambered slides, except that a polyclonal antiserum against *T. pallidum* FlaA protein of (generously provided by Dr. Diane Edmondson) was used as a negative control. Due to the presence of Sf1Ep cells in these slides, images were collected using Nikon Ti2 microscope illuminated using a Lumencor Spectra X light engine and images captured with a Hamamatsu ORCA Flash 4.0 V3 sCMOS camera and Nikon NIS Elements software.

### Opsonophagocytosis Assays

We conducted microscopy-based opsonophagocytosis assay as described before except spirochetes-macrophage coincubation was for 3h (25). In the second approach, J774A.1 macrophages, cultured in DMEM supplemented with 10% FBS, were plated into 24-well, clear-bottom plate with or without coverslips. After washing to remove culture medium, spirochetes were incubated with primary antibodies/normal mouse serum together with pH sensitive pHrodo™ Red, succinimidyl ester in carbonate buffer (prepared to a final concentration of 1 µM) for 1h in the dark, washed and resuspended in PBS/0.2% BSA. Immediately before starting the assay, spirochetes were added to J774A.1 cells and plates were transferred to the IncuCyte ZOOM imaging system maintained at 37 °C with 5% CO₂. An acidic milieu in the late phagosome of macrophages promotes red fluorescence emission by pHRodo dye which can be recorded in real time by the IncuCyte machine (82). Nine images per well were acquired every 30 min for 7h using a 10× objective, with an 800-ms exposure for the red fluorescence channel. Image analysis was performed using IncuCyte Basic Software. Phase-contrast segmentation was achieved by applying a cell-specific mask to exclude background regions, and an area filter was used to eliminate objects smaller than 100 µm². Red-channel background noise was corrected using the Top-Hat method with a 100 µm radius and a threshold of 2 corrected units. The assay wells with coverslips were washed and coverslips were mounted and observed using fluorescence microscope as described above.

### Binding Assays

Binding assays were conducted using B314 expressing TprD/TprK as previously described (81). Three different cell lines were used in this study: HEK293 (a human embryonic kidney epithelial cell line), C6 (a rat glioma line), and HeLa (a human cervical cancer epithelial line). Cell lines were selected to represent tissues colonized by *T. pallidum* during human infection.

### Statistical analysis

All experiments were performed independently at least three times. Statistical significance was assessed for presented representative experiment using an unpaired, two-tailed Student’s *t*-test, with *P* < 0.05 considered statistically significant.

## Authors contributions

NP conceptualized the study, conducted some experiments and finalized the manuscript. KZ and OAZ conducted most experiments, contributed equally to this manuscript while KZ took initiative for different experiments. Both wrote the initial draft of the manuscript. LX and LG infected rabbits and recovered *T. pallidum* for some assays and co-cultured *T. pallidum* with Sf1Ep cells for this study. All authors contributed to editing of the manuscript.

## Acknowledgments

We are grateful to Dr. Diane Edmondson for providing the anti-FlaA antibodies and to Luke Fritzky, manager at the Rutgers Cellular Imaging and Histology Core Facility for taking images of *T. pallidum* after IFA. We also acknowledge Christopher Varsanyi and Luis Carza-Martinez for their assistance with IncuCyte setup and data analysis.

